# Structure and function of accessory Sec proteins involved in the adhesin export pathway of Streptococcus gordonii

**DOI:** 10.1101/219105

**Authors:** Yu Chen, Barbara A. Bensing, Ravin Seepersaud, Wei Mi, Maofu Liao, Philip D. Jeffrey, Asif Shajahan, Roberto N. Sonon, Parastoo Azadi, Paul M. Sullam, Tom A. Rapoport

**Author notes:** Current address: Eisai Inc., 4 Corporate Drive, Andover, MA 01810, USA.

## Abstract

Many pathogenic bacteria, including *Streptococcus gordonii*, possess a pathway for the export of a single serine-rich-repeat protein that mediates the adhesion of bacteria to host cells and the extracellular matrix. These adhesins are *O*-glycosylated by several cytosolic glycosyltransferases and require three accessory Sec proteins (Asp1-3) for export, but how the adhesins are processed for secretion is not well defined. Here, we show that *O*-glycosylation of *S. gordonii* adhesin GspB occurs in a sequential manner by three enzymes (GtfA/B, Nss, Gly) that attach N-acetylglucosamine and glucose to Ser/Thr residues. The modified substrate is subsequently transferred from the last glycosyltransferase to the Asp1/2/3 complex. Crystal structures show that both Asp1 and Asp3 are related to carbohydrate binding proteins. Asp1 also has an affinity for phospholipids, which is attenuated by Asp2. These results suggest a mechanism for the modification of adhesin in the cytosol and its subsequent targeting to the export machinery.

## INTRODUCTION

Adhesion proteins are instrumental for the pathogenicity of bacteria (1). Streptococci and staphylococci bacteria express serine-rich repeat (SRR) adhesins that are exported from the cell, but remain associated with the cell wall and allow the bacteria to attach to the host cells and their extracellular matrix (2, 3). In addition, these adhesins may also mediate interactions between bacteria, facilitating biofilm formation and bacterial colonization (4). The biosynthesis of SRR adhesins is a promising target of novel antibiotics that could be used to treat diseases caused by streptococci and staphylococci, such as infective endocarditis, pneumococcal pneumonia, neonatal sepsis, and meningitis (3).

SRR adhesins use a dedicated pathway for their export from the cytosol, called the accessory Sec system (5, 6); most other proteins are exported from the bacterial cell by the canonical Sec pathway (7). In the canonical pathway, proteins are moved by the SecA ATPase through the protein-conducting SecY channel. In the accessory Sec pathway, export is mediated by distinct SecA and SecY proteins (SecA2 and SecY2). These components are encoded in an operon that also includes the adhesin substrate as well as several glycosyltransferases and accessory Sec system proteins (Asps) (5, 6). The glycosyltransferases attach sugar residues to adhesin before its export from the cytosol (2, 8), but the exact roles of the glycosyltransferases and Asps in the export pathway is not well defined.

The SRR adhesins are initially modified with *N*-acetylglucosamine (GlcNAc) at multiple Ser/Thr residues by the heterodimeric GtfA/B glycosyltransferase (9–14). The deletion of GtfA or GtfB results in non-glycosylated adhesins that are prone to degradation (11, 14, 15). Glycosylation is physiologically important as the deletion of GtfA also reduces the adhesion of bacteria to host cells (15, 16). Recent results show that GtfA is the catalytic subunit, while GtfB is involved in substrate binding (10). Most SRR adhesins are further modified by additional glycosyltransferases that are also encoded by the same operon (5, 6). In *S. parasanguinis* and *S. pneumoniae*, they modify adhesins in a sequential manner (17, 18). In *S. gordonii*, there are two such glycosyltransferases, Nss and Gly (5). Deletion of either enzyme results in compromised modification of the SRR adhesin GspB (9). Nss from related streptococcal species adds glucose to GlcNAc attached to Ser/Thr-containing peptides (19–21). It is unclear how Gly modifies the adhesin GspB, and whether Nss and Gly act sequentially or have redundant functions.

*S. gordonii* encode three Asps (Asp1,2,3), which are conserved among different bacterial species that express SRR adhesins (5, 6). Deletion of any of the Asps blocks the export of the adhesin GspB and results in its intracellular accumulation (15, 22). An essential role for the Asps in the biogenesis of SRR adhesins has also been observed in other species (16, 23–25). Interactions among the Asps and of the Asps with substrate and SecA2 have been reported for both *S. gordonii* and *S. parasanguinis* (where the Asps are called Gaps) (22–24, 26–28), but it remains unclear how the Asps function in GspB export.

Here, we show that the modification of the adhesin GspB of *S. gordonii* by the glycosyltransferases occurs in a sequential manner. First, GlcNAc residues are attached to Ser/Thr residues in the SRR domains of GspB. Next, Nss adds glucose to GlcNAc, and finally, Gly adds glucose to previously attached glucose residues. Interestingly, Gly remains bound to the modified substrate. Release of modified GspB from Gly is caused by the complex of the three Asps (Asp complex). Crystal structures show that indeed both Asp1 and Asp3 are carbohydrate-binding proteins. Asp1 is a catalytically inactive member of the GT-B family of glycosyltransferases and Asp3 contains a carbohydrate-binding module also found in several glycosidases. Our results also show that Asp1 has an affinity for negatively charged phospholipids, which may facilitate substrate delivery to the membrane. Taken together, our results suggest a model for the pathway by which the adhesin is modified and targeted to the export machinery.

## RESULTS

### Glycosyltransferases act in a sequential manner

To test the role of the glycosyltransferases in adhesin modification, we produced a fragment of the GspB substrate by *in vitro* translation in reticulocyte lysate in the presence of ^35^S-methionine. The GspB fragment (GspB-F; Figure S1) contains residues 91 to 736, including the first Ser/Thr-rich domain (SRR1), an intervening sequence that normally binds to host cells (binding region; BR), and the N-terminal part of the second Ser/Thr-rich domain (SRR2N). It lacks the N-terminal signal sequence. GspB-F with the signal sequence is glycosylated in *S. gordonii* cells and secreted with the same efficiency as full-length adhesin (29). *In vitro* translation of GspB-F generated non-glycosylated protein that could be visualized as a single band after SDS-PAGE and autoradiography (Figure 1A; lane 1). As described previously, when a purified complex of GtfA and GftB and UDP-GlcNAc were added after translation, a size shift was observed, caused by modification of GspB-F with GlcNAc residues (G1 species; lane 2). Subsequent addition of purified Nss and UDP-Glc resulted in a further size shift (G2 species; lane 3). Nss did not function with UDP-GlcNAc (Figure 1B; lane 3 versus 2). Finally, yet a larger species was generated when purified Gly was introduced (G3 species; Figure 1A; lane 4). Nss did not attach Glc residues to unmodified GspB-F (Figure 1C; lane 1 versus 3), indicating that it can only modify substrate after GtfA/B has added GlcNAc residues. The same is true for Gly (Figure 1D; lane 1). Finally, modification by Gly was dependent on the prior action of Nss (Figure 1E; lane 3 versus 4). Taken together, these results indicate that GtfA/B, Nss, and Gly function in a defined order; GtfA adds GlcNAc residues to Ser/Thr residues in the SRR domains, which are then further modified with Glc residues by the sequential action of Nss and Gly.

**Figure 1.**
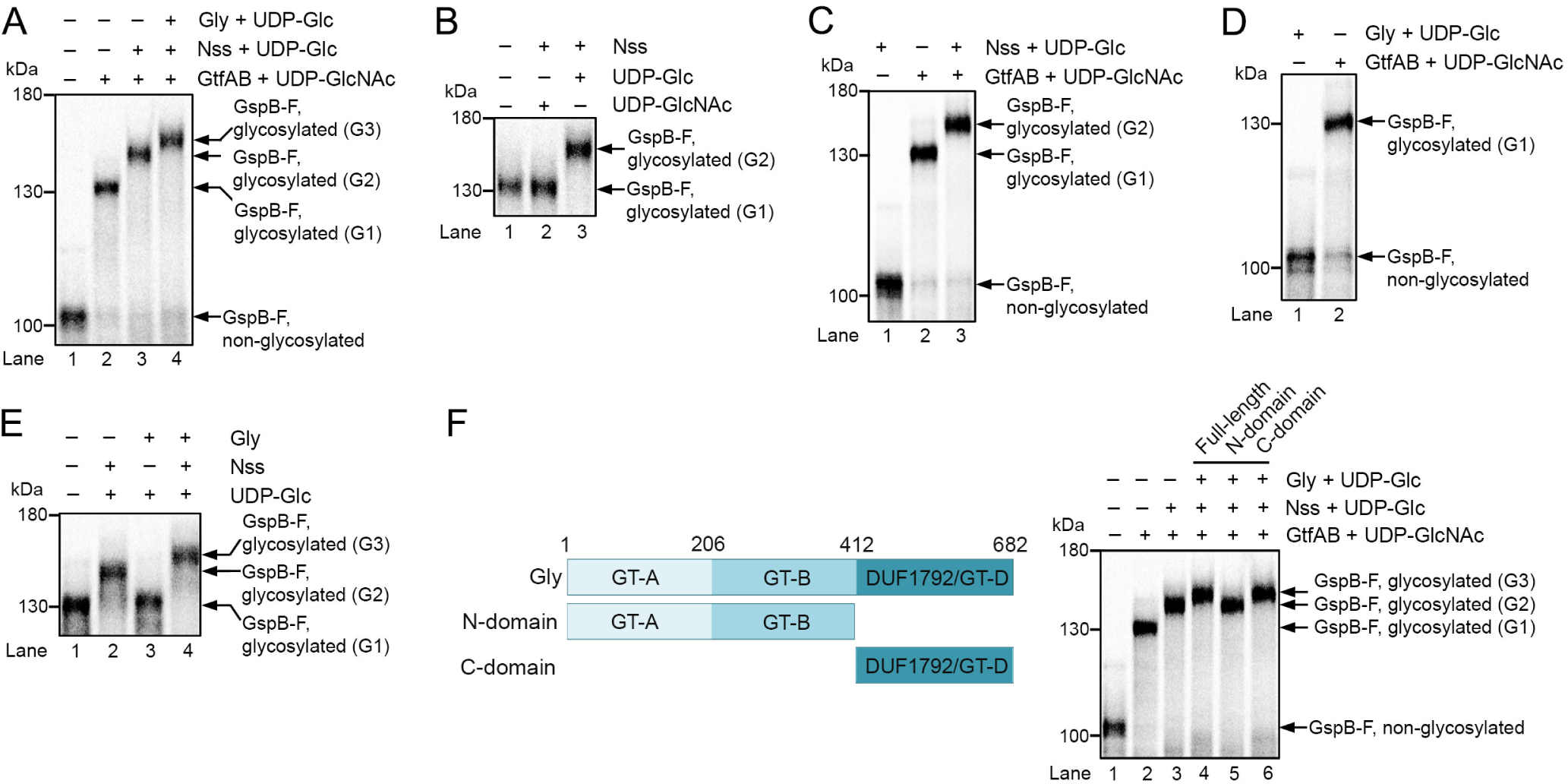
**Adhesin is sequentially *O*-glycosylated by three glycosyltransferases.** (A) A fragment of the *S. gordonii* adhesin GspB (GspB-F) was synthesized in reticylocyte lysate in the presence of ^35^S-methionine. After translation, the glycosyltransferases GtfA/B, Nss, and Gly were added, as indicated, together with UDPGlcNAc or UDP-Glc. The samples were analyzed by SDS-PAGE and autoradiography. G1, G2, and G3 indicate different glycosylated species. (B) *In vitro* synthesized GspB-F was incubated with Nss and either UDP-Glc or UDPGlcNAc. (C) *In vitro* synthesized GspB-F was incubated with UDP-sugars and Nss in the absence or presence of GtfA/B. (D) *In vitro* synthesized GspB-F was modified with GtfA/B or Gly in the presence of UDP-GlcNAc or UDP-Glc, respectively. (E) *In vitro* synthesized GspB-F was modified with GtfA/B and further incubated in the presence of UDP-Glc with either Nss or Gly. (F) The left panel shows the domain organization of Gly. The right panel shows *in vitro* synthesized GspB-F that was incubated with GtfA/B, Nss, and either full-length Gly or its N- or C-terminal domains, together with UDP-Glc or UDP-GlcNAc.

To test whether the modification of GspB with multiple sugars occurs *in vivo*, we purified GspB-F secreted from *S. gordonii* and used mass spectrometry to analyze sugars released by *β*-elimination (Figure S2A). The results show that the protein indeed contains one N-acetyl hexose (HexNAc) and either zero, one, or two hexoses.

Modification with three sugar residues was also seen when GspB-F was expressed in *E. coli* together with GtfA/B, Nss, and Gly (Figure S2B). Although identification of modified GspB-F peptides by mass spectrometry was challenging, we identified with confidence a SRR1 peptide that contained a Ser modified by one GlcNAc and two hexoses (Figure S2C). Taken together, these results are consistent with the idea that Nss and Gly add glucose residues to GlcNAc attached by GtfA/B to Ser/Thr of GspB-F.

Nss consists of a single domain that has a typical GT-B glycosyltransferase fold (20, 21). Gly consists of three domains (Figure 1F; left panel). The first two domains are predicted to have GT-A and GT-B glycosyltransferase folds, respectively, and the third domain has a recently identified GT-D fold (30). Of note, the isolated GT-D domain from a Gly homolog of *S. parasanguinis* has enzymatic activity for its substrate (30). We found that the isolated GT-D of *S. gordonii* Gly was capable of adding Glc residues to GspB-F pre-modified with GtfA/B and Nss (Figure 1F; right panel, lane 6), whereas the isolated N-terminal fragment containing the GT-A and GT-B folds was inactive (lane 5). Thus, despite the fact that the N-terminal domains are sequence-related to glycosyltransferases, they seem to lack enzymatic activity.

### Substrate binds to Gly and is released by the Asp complex

To our surprise, we noticed that fully modified GspB-F remained associated with Gly, the last glycosyltransferase; essentially all G3 species could be recovered with a fusion of Gly with glutathione S-transferase (Gly-GST), followed by binding to a glutathione resin (Figure 2A; lane 2). In contrast, neither GtfA/B (10) nor Nss (lane 1) had appreciable affinity for the product of their modification reactions, as commonly seen for enzymes. The material bound to Gly-GST could be partially released from the beads with a 100-fold excess of full-length Gly (Figure 2B; lane 4), but not with the isolated C-terminal GT-D domain (lane 2). These data suggest that the modified substrate is reversibly bound by the N-terminal GT-A/B domains. This conclusion is supported by the fact that the isolated GT-D domain can generate the G3 species, but does not interact strongly with it, as demonstrated by the absence of competition with the full-length Gly protein (Figure 2C; lane 1). This experiment also shows that the binding of Gly to its enzymatic product is separable from the modification reaction *per se*. In our system, product binding by the N-terminal GT-A/B domains does not interfere with the enzymatic activity of the C-terminal GT-D domain, because Gly is in large excess over substrate and the N-terminal domains binds reversibly to the product.

**Figure 2.**
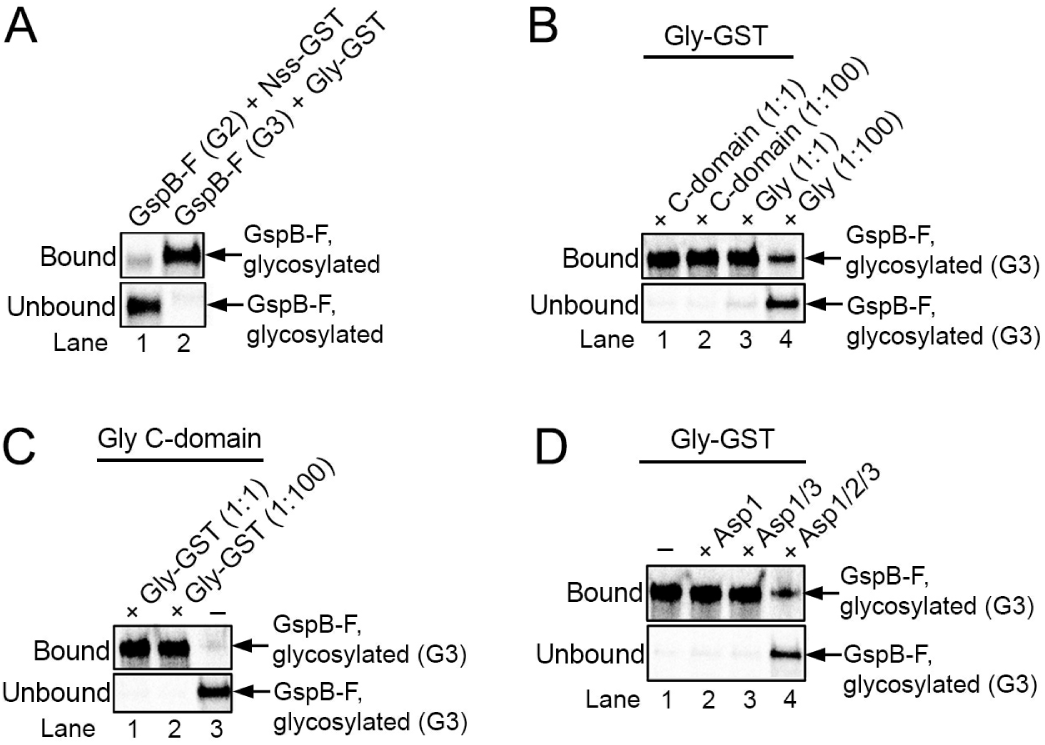
**The glycosyltransferase Gly remains associated with its enzymatic product.** (A) Gsp-F was synthesized *in vitro* in the presence of ^35^S-methionine and then glycosylated with GtfA/B to generate the G1 species, followed by incubation with either a GST-fusion to Nss (Nss-GST; lane 1) or wild type Nss (lane 2) to generate the G2 species. The sample in lane 2 was further incubated with a GST-fusion to Gly (Gly-GST) to generate the G3 species. The samples were then incubated with glutathione-beads, and the bound and unbound fractions analyzed by SDS-PAGE and autoradiography. (B) The G3 species was generated with GtfA/B, Nss, and GST-Gly and bound to glutathione-beads. After washing, the beads were incubated with either the same amount (1:1) or a 100-fold excess (1:100) of His-tagged versions of either full-length Gly (Gly) or C-terminal Gly domain (C-domain). The bound and unbound fractions were analyzed as in (A). (C) The G3 species was generated with a His-tagged version of the C-domain of Gly (Gly C-domain). The sample was then mixed with either the same amount (1:1) or a 100-fold excess (1:100) of Gly-GST and incubated with glutathione-beads. Bound and unbound fractions were analyzed as in (A). (D) The G3 species was generated with Gly-GST and bound to glutathione-beads. After washing, the beads were incubated with Asp1, Asp1/3 complex, or Asp1/2/3 complex, and the bound and unbound fractions were analyzed as in (A).

Next, we tested the role of the Asps. Neither of the three Asps had an effect on the glycosylation reactions catalyzed by GtfA/B, Nss, or Gly (Figure S3). However, the complex of the three Asps released the fully glycosylated G3 species from Gly-GST (Figure 2D; lane 4). Asp1 alone or a complex of Asp1 and Asp3 (Asp1/3) was inactive in the release reaction (lanes 2, 3). These results suggest that the complex of all three Asp proteins may be involved in the transfer of glycosylated substrate from the last glycosyltransferase to the next step in the export pathway.

### Structures of the Asps

Since our data suggest that the Asp complex can accept fully glycosylated substrate from Gly, we suspected that it can interact with carbohydrates. To test this possibility, we determined the crystal structures of Asp1 alone (resolution of 2.77Å) and of an Asp1/3 complex (resolution of 3.11Å) (Table S1). The structure of Asp1 is similar to that of GtfA and GtfB (Figure 3A,B,F). Like GtfA or GtfB, Asp1 has two Rossmann-like folds (R-folds I and II), which are typical for the GT-B family of glycosyltransferases (Figure 3A). In addition, it has the typical extended *β*-sheet domain (EBD). Together, these domains form a U-shaped structure. As in the enzymatically inactive GtfB protein, the cleft between R-folds I and II is negatively charged (Figure 3B). In contrast, GtfA and other enzymatically active GT-B family members have a positively charged cleft that is required to bind UDP-sugars (Figure 3B). Like GtfB, Asp1 lacks two positively charged residues in the active site and has a Gln residue at position 438 in place of an essential Glu residue (Figure 3B, lower panels). The structure thus supports the idea that Asp1, like GtfB, is a carbohydrate binding protein, rather than an active glycosyltransferase. Consistent with the postulated substrate binding site, when two conserved Asp residues in the cleft between the R-folds were mutated to Arg, secretion of GspB-F from *S. gordonii* cells was abolished (Figures 4A, B; sequence alignment shown in Figure S4).

**Figure 3.**
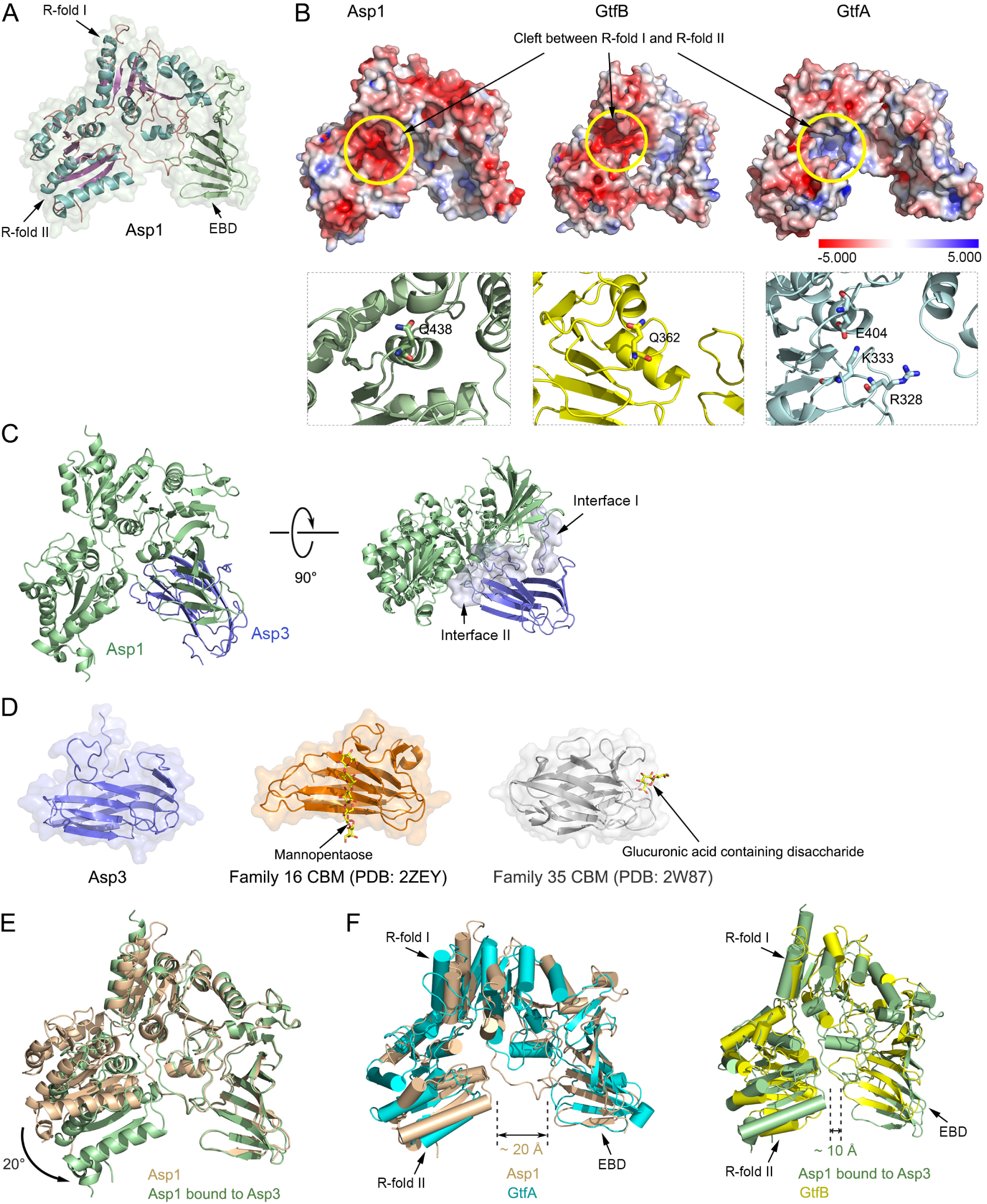
**Crystal structures of Asp1 and Asp3.** (A) Crystal structure of Asp1 alone. Shown is a ribbon diagram of the model imbedded in a space-filling presentation. The helices and the β-sheets of Rossmann folds I and II (R-folds I and II) are shown in cyan and purple, respectively. The extended β-sheet domain (EBD) is shown in green. (B) The upper panels show space-filling models of Asp1, GtfB, and GtfA, with the electrostatic surface calculated with the Adaptive Poisson-Boltzmann Solver, as implemented in Pymol, using a scale from −5.000 to 5.000 (bottom). The yellow circle indicates the cleft between the R-folds, which is negatively charged for Asp1 and GtfB, and positively charged for GtfA. The lower panels show magnified views of the cleft in ribbon presentation. Residues in the active site of GtfA and the corresponding residues in the enzymatically inactive Asp1 and GtfB are shown in stick presentation. (C) Structure of the Asp1/3 complex. Shown is a ribbon diagram of the model, with Asp1 in green and Asp3 in blue. The right panel also shows a space-filling model of the interfaces between Asp1 and Asp3. (D) Comparison of Asp3 with two CBM families. The two CBMs shown bind their sugar substrates at different sites. (E) Comparison of the structures of Asp1 in isolation and when bound to Asp3 (brown and green, respectively). (F) Comparison of the open conformation of Asp1 in isolation with GtfA (left panel; brown and cyan, respectively), and of the closed conformation of Asp1 when bound to Asp3 with GtfB (right panel; green and yellow, respectively).

**Figure 4.**
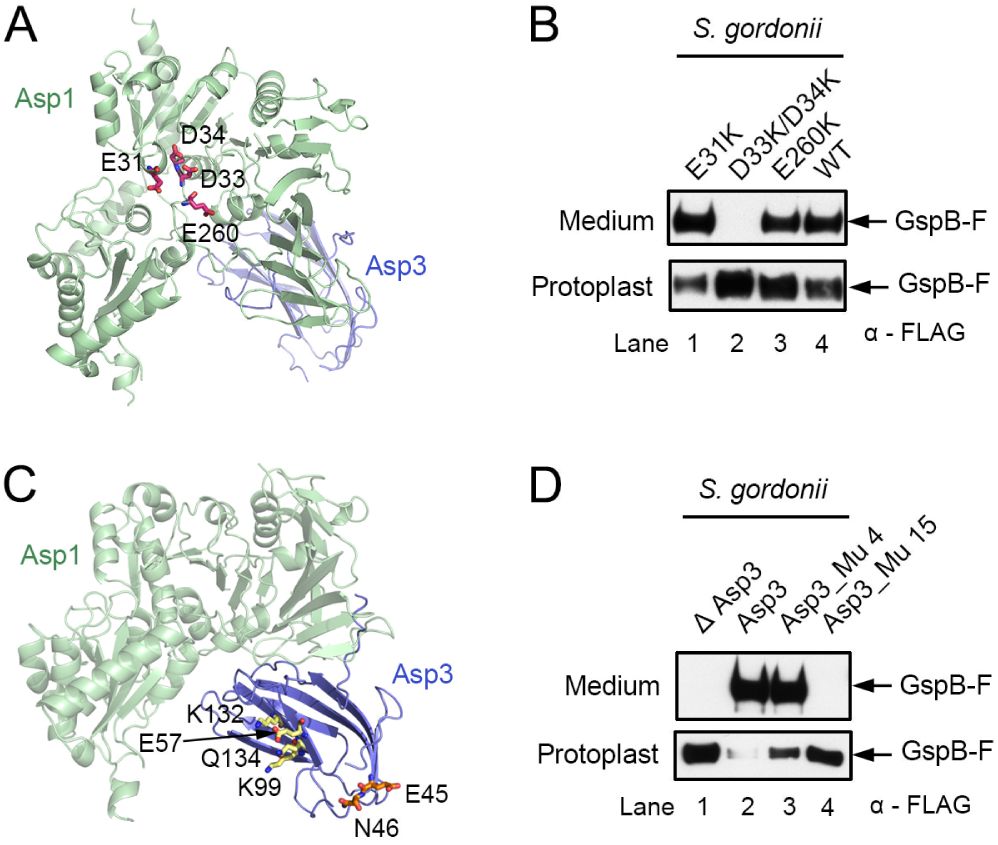
**Mutations in Asp1 and Asp3 affect secretion of adhesin from *S. gordonii* cells.** (A) Ribbon diagram of the Asp1/3 complex (Asp1 in green, Asp3 in blue), with mutated Asp1 residues in the cleft between R-folds I and II shown in stick presentation in magenta. (B) The endogenous Asp1 protein of *S. gordonii* was replaced with the indicated Asp1 mutants. Secretion of FLAG-tagged GspB-F from the cells was tested by subjecting the medium and protoplasts to SDS-PAGE, followed by immoblotting with FLAG antibodies. (C) As in (A), but with mutated Asp3 residues. Mutations in the potential carbohydrate binding regions, the concave surface of the β-sandwich and at the tips of the β-strands, are shown in stick presentation in yellow and orange, respectively. (D) As in (B), but with a *S. gordonii* strain lacking Asp3, expressing either wild type or mutant Asp3. Asp3_Mu4 contains mutations E57K, K132E, K99E, Q134R, and Asp3_Mu15 harbors the mutations E45K and N46K.

Asp1 forms a stable 1:1 complex with Asp3 (Figure S5A). Asp3 could not be stably isolated on its own, suggesting that it has Asp1 as an obligatory partner. Consistent with this observation, deletion of the Asp1 homolog Gap1 in *S. parasanguinis* results in the degradation of the Asp3 homolog Gap3 (28). The Asp1/3 structure shows that Asp3 consists of two anti-parallel *β*-sheets (*β*-sandwich) (Figure 3C). Asp3 uses two different regions to bind to Asp1 (interfaces I and II). Interface I binds to the EBD of Asp1, and interface II to both the EBD and the cleft between the R-folds (Figure 3C). Asp3 is structurally related to carbohydrate binding modules (CBM) in glycosidases (31, 32) (Figure 3D). Interestingly, different CBMs bind their sugar ligands with different surfaces, some with the concave surface of the *β*-sandwich and others with the tips of the *β*-strands (33, 34) (Figure 3D). In the case of Asp3, the latter binding site seems to be more important, as mutations of conserved residues in this area had a drastic effect on GspB-F secretion from *S. gordonii*, whereas mutations in the concave surface of the *β*-sandwich had only a small effect (Figures 4C, D). In the Asp1/3 complex, the Asp1 protein adopts a closed conformation, in which R-fold II moves towards the EBD (Figure 3E). The two ends of the U-shaped structure of Asp1 are much closer in the closed conformation than in the open state (10 Å versus 20 Å; Figure 3F). The open and closed conformations resemble those of GtfA and GtfB, respectively (Figure 3F; ref. (10)). A conformational change from the open to the closed state has been observed for GtfA/B and is likely required for the binding of adhesin.

Asp1, 2, and 3 co-migrated in gel filtration (Figure S5B), but light scattering experiments indicated that there was a mixture of monomeric and dimeric complexes that contain one copy each of Asp1, 2, and 3 (Figure S5C). This heterogeneity is likely the explanation for why these complexes did not crystallize. We therefore attempted to obtain structural information by other means. Addition of trypsin to the Asp1/3 complex generated one Asp1 and one Asp3 peptide, which were not observed with the Asp1/2/3 complex (Figure 5A; indicated by stars). The cleavage sites protected by Asp2 are R430 of Asp1 and R23 of Asp3 (Figure 5B; R23 is in an unstructured region, so the figure shows flanking residues). Thus, Asp2 seems to bind to both Asp1 and Asp3 at the open end of the U-shaped Asp1/3 complex. Next, we used negative-stain electron microscopy (EM) to analyze the Asp1/3 and Asp1/2/3 complexes (Figure 5C, Figure S6). To better locate the individual proteins in the images, we fused the maltose-binding protein (MBP) to Asp1, Asp2, or both. Complexes containing the MBP fusions were monomeric, indicating that the MBP domain interfered with dimerization (Figure S5C). The results confirm that Asp2 sits at the open end of the Asp1/3 complex (Figure 5C, Figure S6). Negative-stain EM also confirmed that without MBP, the Asp1/2/3 complex consisted of a mixture of monomers and dimers. In the dimer, the Asp1/2/3 monomers associate in an anti-parallel fashion (Figure 5C, fourth panel from the left). It is unclear which form of the Asp complex is physiologically relevant.

**Figure 5.**
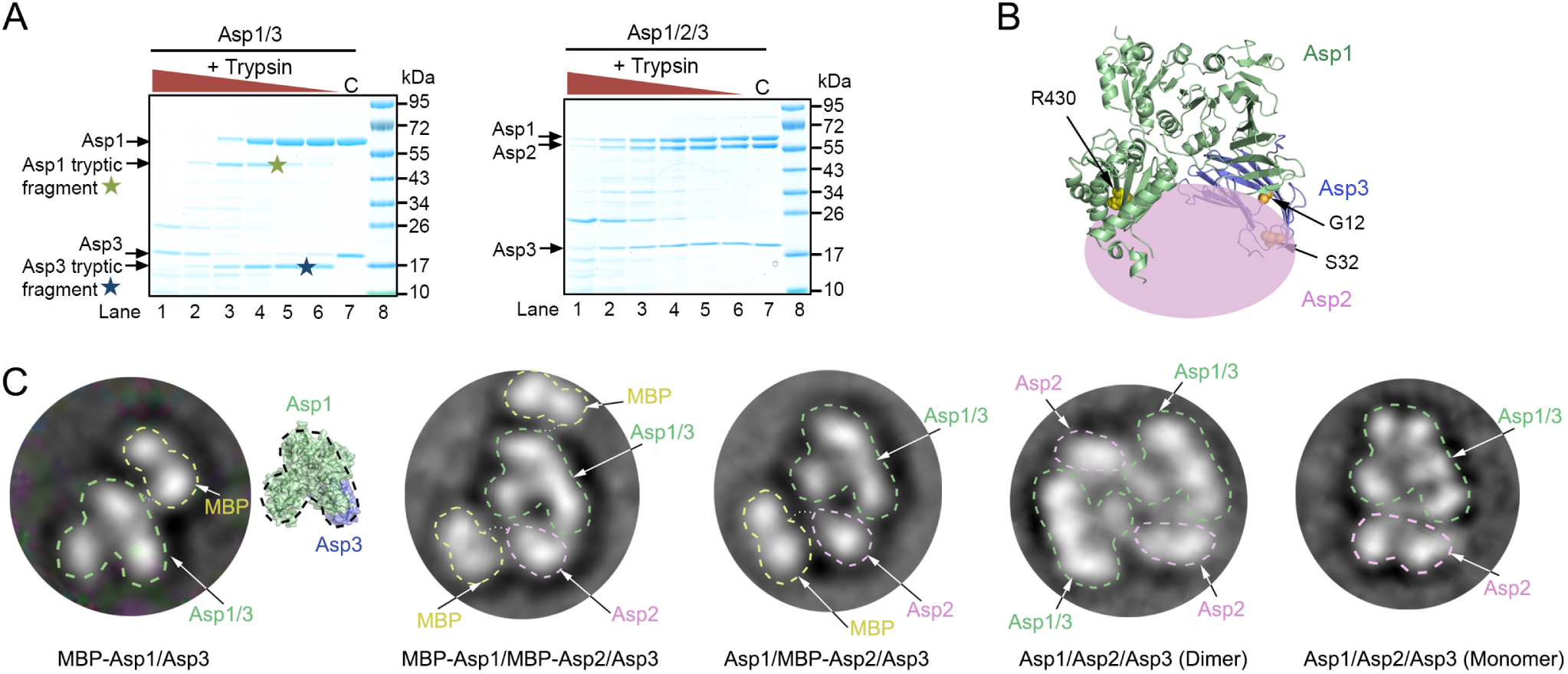
**Domain organization of the Asp1/2/3 complex.** (A) Purified Asp1/3 and Asp1/2/3 complexes were incubated with different concentrations of trypsin. The samples were analyzed by SDS-PAGE and staining with Coomassie blue. Tryptic fragments seen only with Asp1/3 are indicated by stars. C, control without trypsin. (B) Approximate position of Asp2 in the Asp1/2/3 complex. The crystal structure of the Asp1/3 complex is shown as a ribbon diagram and Asp2 as a pink ellipse. Trypsin cleavage at residues R430 and R23 is prevented by Asp2. R430 is indicated as yellow balls. R23 is located in a flexible region that is invisible in the crystal structure, so residues G12 and S32 in the flanking segments are shown (in orange). (C) The left three panels show the domain organization of Asp complexes that contain fusions of maltose binding protein (MBP) to Asp1 and/or Asp2. Shown are 2D averages of particles analyzed by negative-stain electron microscopy. The MBP tag prevents dimerization of the Asp1/2/3 complex. The characteristic two-globule shape of MBP and the domain structure of Asp1/3, as deduced from a comparison with the crystal structure, were used to identify the proteins. The right two panels show the domain organization of dimeric and monomeric untagged Asp1/2/3 complex.

### Substrate targeting to membranes by Asp1/3

After being released from Gly by the Asp complex, the glycosylated substrate needs to be targeted to the membrane, a process that might be mediated by the Asps. We therefore tested whether the Asps have an affinity for membranes. To this end, purified Asps were incubated with liposomes of different phospholipid composition. The samples were then subjected to flotation in a Nycodenz gradient and fractions were analyzed by SDS-PAGE and Coomassie staining (Figure 6A). With liposomes containing a high percentage of negatively charged lipids (dioleoylphosphatidylglycerol; DOPG), Asp1 alone or Asp1/3 floated to the second fraction from the top (Figure 6B), which is also the peak position of the lipids (Figure S7). The Asp1/2/3 complex also bound to the liposomes, but it peaked at fraction 3, suggesting that Asp2 weakens the interaction with the liposomes. Indeed, when the percentage of negatively charged lipids was decreased, the binding of Asp1/2/3 was selectively reduced (Figures 6C, D). These results indicate that Asp1 and Asp1/3 have an affinity for negatively charged lipids. Given that Asp1 and Asp3 are always in a complex, Asp1 is likely responsible for membrane targeting of both proteins. Asp2 inhibits membrane interaction, suggesting that lipid head groups and Asp2 may compete for interaction with the Asp1/3 complex. No interaction of Asp1 and Asp1/3 with polar lipids from *E. coli* was observed (Figure 6E), consistent with the fact that *E. coli* contains a much lower percentage of negatively charged lipids than do streptococci (35–37).

**Figure 6.**
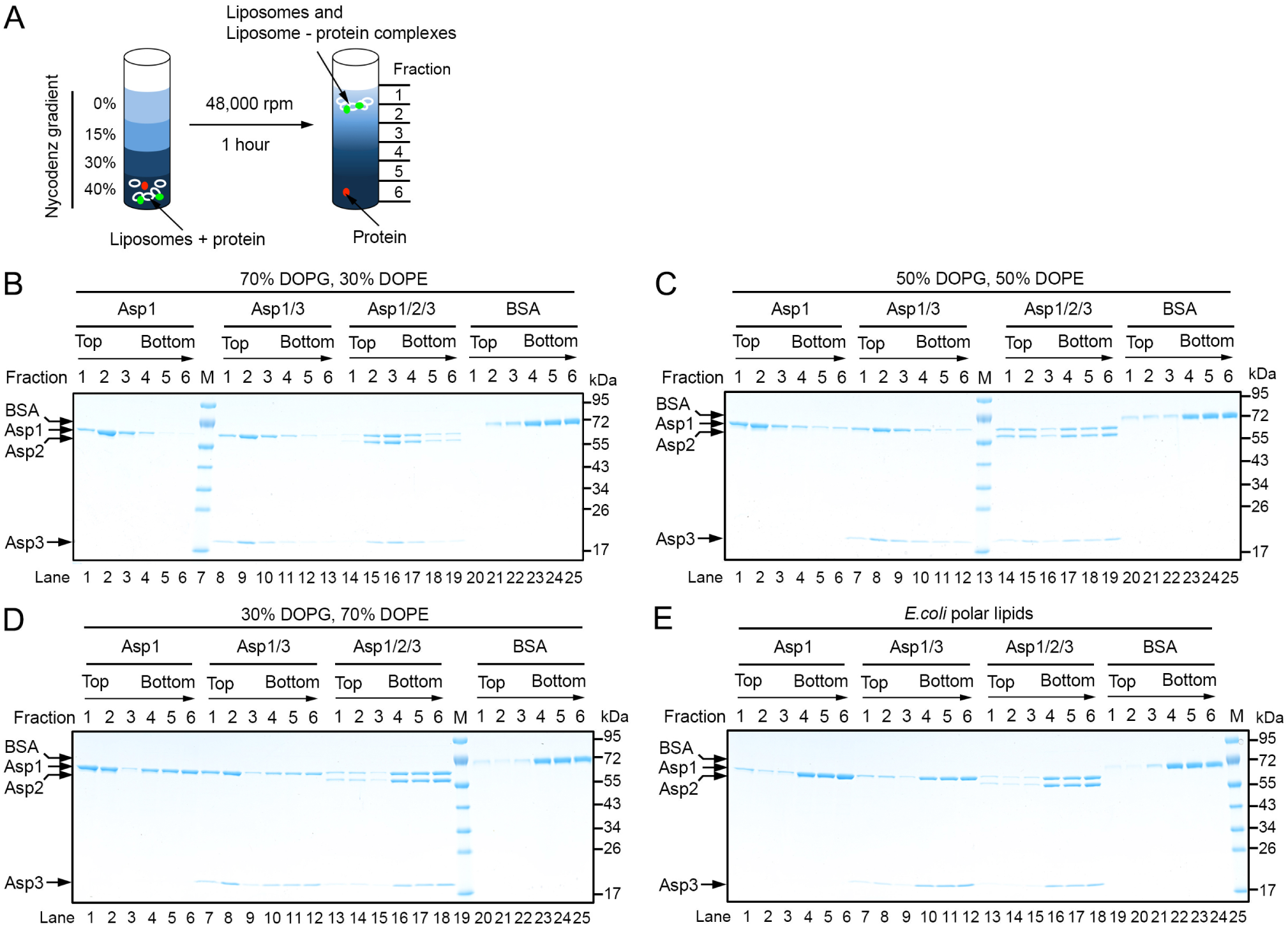
**Interaction of the Asps with phospholipid bilayers.** (A) Scheme of the binding assay. Liposomes containing different lipid compositions were mixed with Asps and the samples were subjected to flotation in a Nycodenz gradient. Fractions were collected from the top and analyzed by Coomassie staining. Liposome-bound proteins (green) are expected to co-float with the lipids (white). Non-associated proteins (red) stay at the bottom. (B)-(D) The indicated combinations of Asps were tested for binding to liposomes containing a different ratio of the negatively charged lipid dioleoylphosphatidyl glycerol (DOPG) and the neutral lipid dioleoylphosphatidyl ethanolamine (DOPE). Lipids peak in fraction 2. Bovine serum albumin (BSA) was used as a control. (E) Binding of the Asps to liposomes generated with *E. coli* polar lipids.

To test whether a glycosylated substrate can be targeted to the membrane by the Asps, we incubated glycosylated GspB-F with either Asp1, Asp1/3, or Asp1/2/3, followed by incubation with liposomes containing negatively charged lipids. The vesicles were then floated in a Nycodenz gradient and fractions analyzed by immunoblotting for a FLAG tag on GspB-F and Coomassie staining. With either Asp1 or Asp1/3, a small, but reproducible fraction of the substrate floated with the liposomes (Figure 7A). No coflotation was observed with Asp1/2/3 or without the Asps. An Asp1/3 complex containing Asp3 mutations at the tip of the *β*-strands, which abolished GspB-F secretion *in vivo* (Figure 4C,D; mutant 15), was almost completely defective in substrate flotation (Figure 7B). Asp3 mutations in the concave surface area had a more moderate effect, again consistent with *in vivo* results. These results show that a substrate can be recruited by the Asp1/3 complex to the membrane.

**Figure 7.**
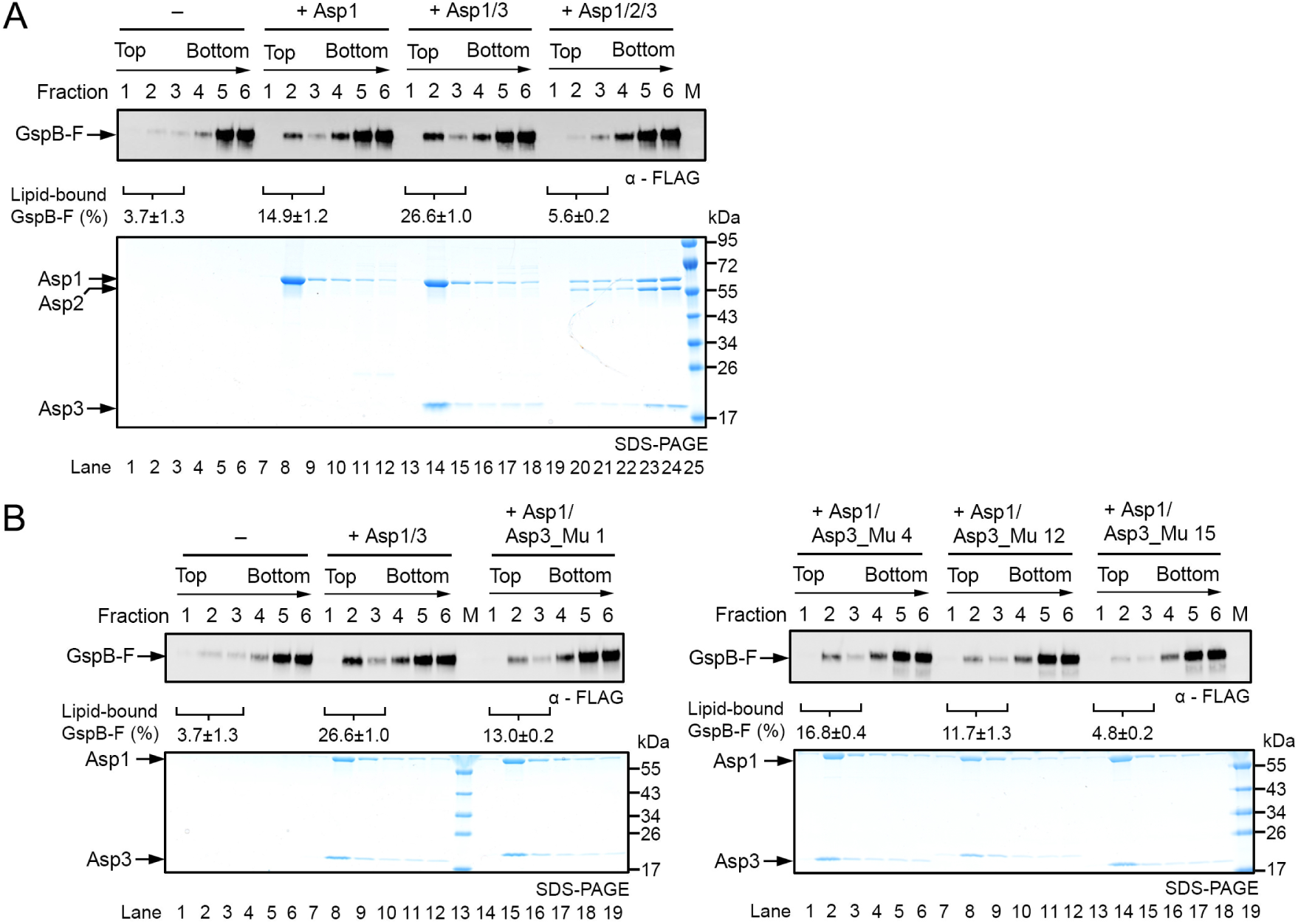
**Asps target adhesin to the membrane.** (A) Glycosylated, FLAG-tagged GspB-F was purified from *S. gordonii* cells lacking Nss and Gly, but the glycans contain significant amounts of hexose attached to GlcNAc, probably because of other glycosyl transferases. The purified protein was incubated with different combinations of the Asps, as indicated, followed by incubation with liposomes containing 70% DOPG, 29.5% DOPE and 0.5% Texas Red-DHPE. The liposomes were floated in a Nycodenz gradient and fractions were analyzed by SDS-PAGE, followed by immunoblotting with FLAG antibodies and Coomassie blue staining (upper and lower panels, respectively). The percentage of membrane-associated Gsp-B was quantified in three experiments and is given underneath the FLAG immunoblot as means and standard error of the means. (B) As in (A), but with Asp1/3 complexes containing either wild type or mutant Asp3. Asp3_Mu 1: E57K, K132E, Q134R; Asp3_Mu 4: E57K, K132E, K99E, Q134R; Asp3_Mu 12: K132E; Asp3_Mu 15: E45K, N46K. The quantification with the mutant complexes shows the means and standard error of two experiments.

## DISCUSSION

Our results suggest a model for the first steps in the export of an SRR adhesin from the pathogenic bacterium *S. gordonii*. The adhesin (GspB) is first made as an unmodified protein. It is then sequentially glycosylated by three glycosyltransferases (see model in Figure 8, box 1). The first enzyme, GftA/B adds GlcNAc residues to Ser/Thr residues in SRR domains (G1 species). Next, Nss adds Glc to the GlcNAc residues (G2 species), and finally, Gly adds further Glc residues to those attached by Nss (G3 species). The fully glycosylated substrate is then transferred to the Asp1/2/3 complex (box 2). In the next step, the Asp1 protein would mediate the interaction of the Asp1/2/3 complex with the lipid bilayer (box 2). The Asp1/2/3 complex has a relatively low affinity for the membrane, so we assume that it continuously cycles between the cytosol and membrane, with the majority staying in the cytosol. Membrane binding of the Asp complex probably requires a conformational change to expose a lipid interaction domain on Asp1, an interface that seems to be fully available for membrane interaction in Asp1 or the Asp1/3 complex. Once at the membrane, the substrate would be delivered to SecA2 and SecY2 for translocation across the membrane (box 3). This is consistent with previous findings that the Asp1/2/3 localizes near SecA2 at the membrane (38).

**Figure 8.**
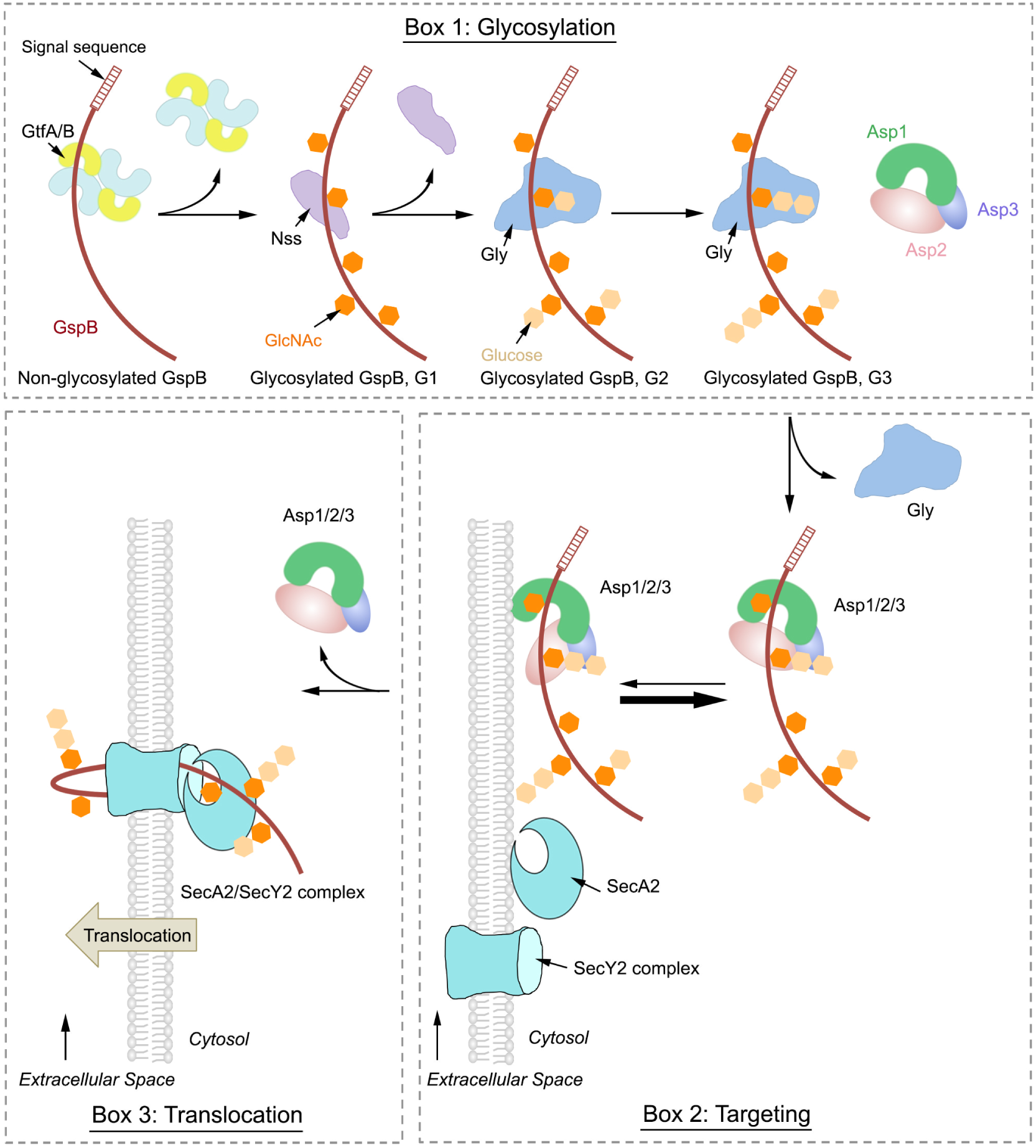
**Model for the export of adhesin from S. gordonii.** Box 1: The adhesin GspB is synthesized and then sequentially *O*-glycosylated by three glycosyltransferases. The first enzyme is GtfA/B, a tetramer that adds GlcNAc residues to Ser/Thr residues (G1 species). Then, Nss adds Glc to GlcNAc (G2 species), and finally Gly adds Glc to Glc residues (G3 species). Gly remains bound to the G3 species, until it is transferred to the Asp1/2/3 complex. Box 2: Although the Asp complex has a low affinity for the lipid bilayer, it probably continuously cycles between the cytosol and membrane. The Asp complex likely undergoes a conformational change, in which Asp2 moves away from a lipid binding surface, allowing the Asp complex to deliver the substrate GspB to the membrane. Box 3: GspB engages SecA2 and the SecY2 channel for its translocation across the membrane and the Asp complex returns to the cytosol.

Our data show that the glycosyltransferases act in a strictly sequential manner. GtfA/B only modifies Ser/Thr residues and has specificity for GlcNAc, while Nss and Gly recognize GlcNAc and Glc residues, respectively. Gly is an unusual enzyme, as it has affinity for the product of the reaction it catalyzes. This is explained by the fact that the enzymatic reaction is catalyzed by the C-terminal GT-D domain, whereas the binding to the product is mediated by the N-terminal GT-A and -B domains. Despite their similarity with glycosyltransferases, the GT-A and –B domains seem to belong to the class of catalytically inactive carbohydrate binding proteins, which also include GtfB and Asp1. Although the affinity of the GT-A and -B domains is high, glycosylated substrate can still dissociate from Gly, allowing its binding to the downstream Asp complex.

Surprisingly, both Asp1 and Asp3 are structurally related to carbohydrate-binding proteins. Asp1 is a catalytically inactive member of the GT-B family of glycosyltransferases and Asp3 is similar to the CBM domain of glycosidases. The presence of carbohydrate-binding motifs in Asp1 and Asp3 strongly supports the idea that they bind glycosylated adhesin. Indeed, our mutagenesis data provide evidence for substrate interaction with Asp3. Asp1 also seems to bind substrate, as it allows co-flotation with liposomes. The interactions of Asp1 and Asp3 with substrate are weak, as they do not survive in pull-down or gel filtration experiments (data not shown). However, substrate release from Gly (Fig. 2) and our co-flotation experiments (Fig. 7) indicate that the Asps do interact with substrate. The low binding affinity is in fact typical for carbohydrate-binding proteins (39). Our results do not exclude the possibility that, under certain conditions, the Asps can also interact with non-glycosylated adhesin. In fact, Asp1 has a similar structure as GtfB (Figure 3B), which can bind non-glycosylated substrate (10), and Asp2 and Asp3 have been shown to bind non-glycosylated GspB (40). Such an interaction would explain why GspB is secreted in an Asp-dependent manner in glycosylation-defective *S. gordonii* strains, although in this situation, much of the substrate is degraded (22). It should also be noted that some bacterial species export SRR adhesins in an Asp-dependent manner although they lack Gly and Nss (5). Asp1 also has an affinity for the lipid bilayer, which facilitates the recruitment of substrate to the membrane. Given that binding of Asp1 requires negatively charged phospholipids and is enhanced at higher salt concentrations (compare Figure 6B (100 mM) with Figure 7A (300 mM)), it seems that it is mediated by both hydrophobic and electrostatic interactions. Taken together, our data indicate that Asp1 and 3 are carbohydrate- and lipid-binding proteins. We favor a model in which the Asps facilitate the transfer of glycosylated substrate to the membrane (Figure 8), rather than simply prevent their premature folding in the cytosol; the repetitive structure of the SRR domains and their extensive *O*-glycosylation would prevent the folding or aggregation of substrate even in the absence of the Asps. Indeed, under these conditions, glycosylated GspB accumulates in the cytosol as soluble protein (15).

How the substrate is delivered to SecA2 and SecY2 remains to be clarified. The signal sequence of adhesin and the adjacent “accessory Sec transport” (AST) domain are required to target the precursor to the accessory Sec system and initiate translocation (41). Although not very hydrophobic (5), the signal sequence could still facilitate the interaction with the lipid bilayer. Once the substrate is bound to the membrane, it could associate with SecA2, a transfer that may be facilitated by an interaction between the Asp complex and SecA2 (22, 38). Based on a homology model, SecA2 has a pronounced positively charged surface patch (Figure S8A, B), which could mediate its interaction with negatively charged phospholipids in the membrane. No such basic surface patch is seen in a homology model for *S. gordonii* SecA1 (Figure S8C). Since we were unsuccessful in purifying soluble SecA2, even in the presence of detergent, we speculate that SecA2 requires a lipid environment to maintain its native conformation and that it is permanently bound to the membrane, in contrast to SecA1, which cycles between the cytosol and membrane (7). According to the model, SecA2 would rely on the Asps to deliver substrate to the membrane where SecA2 could engage both the AST domain and the remainder of the mature domain, whereas in the canonical secretion system, SecA1 would do the job, with the chaperone SecB acting upstream for some substrates and bacteria (7). Once substrate has been recruited to the SecA2/SecY2 complex, it is likely translocated across the membrane by a mechanism similar to that of the canonical system.

## EXPERIMENTAL PROCEDURES

Details are given in the Supporting Information.

### Purification of proteins

All proteins were expressed in *E. coli.* GtfA/B complex was prepared as previously described (10). *S. gordonii* Nss, Gly, Gly N-domain (residue 1-411), and Gly C-domain (residue 412 - 682) were purified utilizing a His-tag. Nss and Gly were also expressed with a C-terminal glutathione S-transferase (GST) tag. Asp1 was expressed with a C-terminal GST tag, either alone or together with Asp3. Purification was performed with glutathione sepharose 4B beads, the GST portion was cleaved off, and the proteins were further purified by ion exchange and gel filtration chromatography. Mutations were introduced into Asp1 or Asp3 by QuikChange mutagenesis. For electron microscopy, untagged Asp1 and Asp3 were co-expressed with Asp2 fused its N-terminus with either maltose binding protein or GST. After purification on either amylose or glutathione resin, the proteins were further purified by ion exchange chromatography and gel filtration. To generate the Asp1/2/3 complex without a tag, the GST tag was cleaved from Asp2 by thrombin protease, and subsequently removed by gel filtration.

### *In vitro* glycosylation assays

GspB-F was generated by *in vitro* translation in reticulocyte lysate in the presence of ^35^S-methionine and analyzed as previously described (10). *In vitro* glycosylation was performed at 37 °C. GtfA/B (1.6 μM), 10 mM UDP-GlcNAc, and GspB-F (2 μl) were first incubated in 5 μl for 10 min, followed by 6.4 μM Nss and 10 mM UDP-glc for 20 min, and 6 μM Gly and 10 mM UDP-Glc for 1 hr. Where indicated, Asp1, Asp1/3, or Asp1/2/3 were included in the reaction at 17 μM.

### Pull-down experiments

*In vitro*-synthesized substrate binding of Nss and Gly was tested with GST-tagged enzymes and magnetic glutathione resin. Where indicated, Asp proteins, Gly, or Gly-C domain were added prior to the resin. Bound and unbound fractions were analyzed by SDS-PAGE and autoradiography.

### Crystallization and structure determination

Crystallization of Asp1 and Asp1/3 complex was performed by the hanging-drop vapor-diffusion method at 22 °C. Both native and selenium SAD data sets were collected at beamline 24ID-C at the Argonne National Laboratory and processed with XDS (42) and autoPROC (43). The positions of Se atoms were determined and phases were calculated using AutoSol Wizard in PHENIX (44). A complete model of Asp1/3 was built in Coot (45). The structure was refined with Phenix.refine (46). The structure of Asp1 was determined by molecular replacement using PHASER in PHENIX (44), with the Asp1 lacking the R-fold II as the initial search model. The model was modified in Coot (45) and refined with Phenix.refine (46). Figures showing structures were prepared in PyMOL (Version 1.5.0.4 Schrödinger, LLC.). All software packages were accessed through SBGrid (47).

### Secretion of GspB-F from *S. gordonii*

Asp1 or Asp3 variants were cloned into plasmid pMSP3545 as described (15) and introduced into *S. gordonii* strain PS1242 (gspB::pB736flagC ∆asp1::spec) or PS1244 (gspB::pB736flagC ∆asp3::spec), respectively (22, 38), by natural transformation (48). GspB-F secretion was determined by immunoblotting with anti-FLAG monoclonal antibodies (Sigma) as described before (29).

Secreted GspB-F was purified from a *S. gordonii* carrying signal sequence-containing GspB-F in place of wild type GspB. GspB-F was enriched from the medium by ammonium sulfate precipitation. After dialysis, glycosylated the protein was purified with a resin containing succinylated wheat germ agglutinin (sWGA), followed by gel filtration.

### Negative stain electron microscopy

Negatively stained specimens were prepared as descibed (49). Grids were imaged on a Tecnai T12 electron microscope (FEI) operated at 120 kV at a nominal magnification of 67,000x using a 4k x 4k CCD camera (UltraScan 4000, Gatan), corresponding to a calibrated pixel size of 1.68 Å on the specimen level. The images were processed as described (50).

### Limited trypsin proteolysis

Asp1/3 or Asp1/2/3 (5.6 μg) were incubated with 6.6, 2.2, 0.73, 0.24, 0.08, or 0.03 μg trypsin protease in 8 μl buffer containing 20 mM Tris/HCl, pH 8.0, and 100 mM NaCl at 22 °C for 20 min. Tryptic fragments of Asp1 and Asp3 were subjected to analysis by MALDI-TOF mass spectrometry (Taplin Mass Spectrometry Core Facility, Harvard Medical School) and N-terminal sequencing (Tufts University Core Facility).

### Liposome flotation assay

Liposomes were generated with either dioleoylphosphatidyl glycerol (DOPG) and dioleoylphosphatidyl ethanolamine (DOPE) or with *E. coli* polar lipids (Avanti). Texas Red-DHPE (ThermoFisher Scientific) was included in some experiments. Asp1, Asp1/3, Asp1/2/3, and BSA were mixed with liposomes at a 1:1,300 molar ratio of protein to lipid and floated in a discontinuous 0-40% (w/v) Nycodenz gradient prepared in 20 mM Tris/HCl, pH 7.5, 100 mM NaCl, in a TLS-55 swinging bucket rotor (Beckman Coulter) at 48,000 rpm for 1 hr. Fractions were collected from the top of the gradient and analyzed by SDS-PAGE and Coomassie blue staining.

Glycosylated GspB-F was isolated from *S. gordonii* cells as described before (10) and incubated with Asps prior to the addition of liposomes. The Nycodenz gradient was prepared in 20 mM Tris/HCl, pH 7.5, 300 mM NaCl. After centrifugation, fractions were subjected to SDS-PAGE and immunoblotting using Flag antibodies (Sigma).

### Glycan and glycol-peptide analysis by Mass spectrometry

Glycosylated GspB-F was subjected to SDS-PAGE. Gel slices were treated with trypsin, and glycopeptides were analyzed on an Orbitrap Fusion mass spectrometer equipped with a nanospray ion source connected to a Dionex binary solvent system. The LC-MS/MS spectra of tryptic peptides of glycosylated GspB-F were searched against the sequence of GspB-F using Proteome Discoverer 1.4 software.

For glycan analysis, *O*-glycans were *β*-eliminated by treatment of extracted tryptic peptides with NaOH/NaBH_4_. The glycans were permethylated and subjected to ESI-MS/MS analysis on an Orbitrap Fusion mass spectrometer. For monosaccharide analysis, tryptic peptides were hydrolyzed with trifluoroacetic acid.

### Homology modeling

Models of *S. gordonii* SecA1 and SecA2 were generated by SWISS-MODEL (51–54), using *T. maritima* SecA (PDB: 3DIN) (55) and *B. subtillis* SecA (PDB: 3JV2) (56) as templates, respectively. The predicted models have GMQE scores of 0.69 and 0.72, respectively.

## ACKNOWLEDGEMENTS

We thank the staff at the Advanced Photon Source of the Northeastern Collaborative Access Team (NE-CAT) beamline for help with data collection. NE-CAT is supported by a grant from the National Institute of General Medical Sciences (P41 GM103403) from the National Institutes of Health (NIH). The Pilatus 6M detector on 24-ID-C beam line is funded by a NIH-ORIP HEI grant (S10 RR029205). This research used resources of the Advanced Photon Source, a U.S. Department of Energy (DOE) Office of Science User Facility operated for the DOE Office of Science by Argonne National Laboratory under Contract No. DE-AC02-06CH11357. Y.C. was supported by an HHMI-Helen Hay Whitney Foundation fellowship. The work in the laboratory of T.A.R. was supported by NIH grant GM052586. T.A.R. is a Howard Hughes Medical Institute Investigator. P.M.S. is supported by the Department of Veterans Affairs, NIH grants R01-AI041513 and R01-AI106987, and the Northern California Institute for Research and Education. P.A. is supported by the NIH-funded Research Resource for Integrated Glycotechnology (NIH grants 1S10OD018530 and P41GM10349010) and U.S. Department of Energy grant DE-FG02-93ER20097 to the Complex Carbohydrate Research Center.

## Author contributions

Y.C. and T.A.R. designed research; Y.C., B.A.B., W.M., M.L., P.D.J, A.S., R.N.S. performed research; Y.C., B.A.B, R.S., P.M.S., M.L., P.A., and T.A.R. analyzed data; and Y.C. and T.A.R. wrote the paper.

## Data deposition

Crystallography, atomic coordinates, and structure factors have been deposited in the Protein Data Bank, www.pdb.org (PDB ID codes 5VAE and 5VAF).

